# A genome resequencing-based genetic map reveals the recombination landscape of an outbred parasitic nematode in the presence of polyploidy and polyandry

**DOI:** 10.1101/177550

**Authors:** Stephen R. Doyle, Roz Laing, David J. Bartley, Collette Britton, Umer Chaudhry, John S. Gilleard, Nancy Holroyd, Barbara K. Mable, Kirsty Maitland, Alison A. Morrison, Andy Tait, Alan Tracey, Matthew Berriman, Eileen Devaney, James A. Cotton, Neil D. Sargison

## Abstract

The parasitic nematode *Haemonchus contortus* is an economically and clinically important pathogen of small ruminants, and a model system for understanding the mechanisms and evolution of traits such as anthelmintic resistance. Anthelmintic resistance is widespread and is a major threat to the sustainability of livestock agriculture globally; however, little is known about the genome architecture and parameters such as recombination that will ultimately influence the rate at which resistance may evolve and spread. Here we performed a genetic cross between two divergent strains of *H. contortus*, and subsequently used whole-genome re-sequencing of a female worm and her brood to identify the distribution of genome-wide variation that characterises these strains. Using a novel bioinformatic approach to identify variants that segregate as expected in a pseudo-testcross, we characterised linkage groups and estimated genetic distances between markers to generate a chromosome-scale F_1_ genetic map composed of 1,618 SNPs. We exploited this map to reveal the recombination landscape, the first for any parasitic helminth species, demonstrating extensive variation in recombination rate within and between chromosomes. Analyses of these data also revealed the extent of polyandry, whereby at least eight males were found to have contributed to the genetic variation of the progeny analysed. Triploid offspring were also identified, which we hypothesise are the result of nondisjunction during female meiosis or polyspermy. These results expand our knowledge of the genetics of parasitic helminths and the unusual life-history of *H. contortus,* and will enable more precise characterisation of the evolution and inheritance of genetic traits such as anthelmintic resistance. This study also demonstrates the feasibility of whole-genome resequencing data to directly construct a genetic map in a single generation cross from a non-inbred non-model organism with a complex lifecycle.

**Author summary:** Recombination is a key genetic process, responsible for the generation of novel genotypes and subsequent phenotypic variation as a result of crossing over between homologous chromosomes. Populations of strongylid nematodes, such as the gastrointestinal parasites that infect livestock and humans, are genetically very diverse, but little is known about patterns of recombination across the genome and how this may contribute to the genetics and evolution of these pathogens. In this study, we performed a genetic cross to quantify recombination in the barber’s pole worm, *Haemonchus contortus*, an important parasite of sheep and goats. The reproductive traits of this worm make standard genetic crosses challenging, but by generating whole-genome sequence data from a female worm and her offspring, we identified genetic variants that act as though they come from a single mating cross, allowing the use of standard statistical approaches to build a genetic map and explore the distribution and rates of recombination throughout the genome. A number of genetic signatures associated with *H. contortus* life history traits were revealed in this analysis: we extend our understanding of multiple paternity (polyandry) in this species, and provide evidence and explanation for sporadic increases in chromosome complements (polyploidy) among the progeny. The resulting genetic map will aid in population genomic studies in general and enhance ongoing efforts to understand the genetic basis of resistance to the drugs used to control these worms, as well as for related species that infect humans throughout the world.

## Background

Recombination is a key genetic process: the breaking and re-joining of genetic material to produce novel genotypes and in turn, generate phenotypic variation. In eukaryotes, this is achieved by crossing-over between homologous chromosomes during the generation of gametes in meiosis. A common approach to studying recombination is to perform controlled matings (i.e. genetic crosses) between genetically distinct and inbred parents. The parents and offspring are then genotyped to construct genetic linkage maps, which aim to order genes or genetic markers based on the recombination frequency between them. This approach can also be used to identify regions of the genome underlying phenotypic variation, and has been widely used for mapping both simple and complex traits in a range of different organisms [1, 2]. More recently, as whole-genome sequencing data has become available for many organisms, genetic maps have been used to inform or validate contig order in genome assemblies [3-7]. Where a contiguous genome assembly is already available, a linkage map can be used to explore variation in recombination rates throughout the genome [8] and determine how this has shaped other aspects of genome architecture, such as the distribution of repeats or the impact of natural selection.

Understanding variation in the rate and pattern of recombination is critical, both for designing and analysing experiments aimed at mapping the genetic basis of phenotypic traits and in interpreting genetic variation in natural populations. Between species, a negative relationship between genome size and recombination rate has been described [9]. Within a species, variation in recombination rate is strongly influenced by the sex of the organism; recombination may not occur in one of the two sexes (typically the heterogametic sex, i.e. the Haldane-Huxley rule [10]), or, if recombination does occur in both sexes, then females tend to show a higher recombination rate than males (i.e. heterochiasmy [11]). In addition, recombination rates have been show to vary considerably within and between chromosomes, which has been attributed to genomic features including but not limited to GC content, gene density, gene size, simple repeats, and chromatin state [12-15]. Among nematodes, recombination is best characterised in the model organism *Caenorhabditis elegans*, where direct comparison of the physical and genetic maps clearly reveals asymmetrically distributed high and low recombination rate domains in each chromosome, correlated with low and high gene density (and gene expression), respectively [8, 14, 16]. However, the precise local DNA features that mediate these rate changes remain unclear. Even less is known about recombination in parasitic helminths. Low density genetic maps are available for only three species, the root knot nematode *Meloidogyne hapla* [5, 6], the human blood-fluke *Schistosoma mansoni* [17], and the rat gastrointestinal parasite *Strongyloides ratti* [7], and only discrete regions of recombination variation have been described in *M. hapla* [5]. Recombination rate variation has been proposed to influence the distribution of genetic variation, and in turn, evolution of phenotypic traits in *C. elegans* [18-20]. Therefore, understanding genome-wide recombination variation in parasitic species will likely be important in predicting the genetic architecture and evolution of important parasite life history traits, including pathogenicity, response to host immunity and chemotherapeutic selection.

The parasite *Haemonchus contortus* is amongst the most pathogenic of the gastrointestinal nematodes and exerts significant burdens on animal health and the economic viability of livestock farming [21]. It is also an emerging model for the biology of parasitic helminths more widely, particularly for understanding anthelmintic drug action and resistance [22]. In particular, *H. contortus* is the most genetically tractable of any of the strongylid (clade V) parasitic nematodes, a large and important group of parasites including key human and veterinary pathogens. It makes a particularly good model because: (i) it is a sexually reproducing diploid organism for which the karyotype—five autosomes and XX/XO sex chromosomes—is well defined [23]; (ii) two published draft genome sequences and extensive transcriptomic data are available [24-26]; (iii) it is amenable to cryopreservation of isolates; and (iv) it is one of the few parasitic nematode species in which genetic crosses have been successfully established [27-33].

Anthelmintic drug failure is an important economic and animal health problem, as anthelmintic resistance is widespread on farms, and populations and isolates resistant to all major classes of anthelmintics have been described [34-36]. Accordingly, significant research effort is focused on the development of novel anthelmintics [37] or vaccines [38] for parasite control. Although research on *H. contortus* has been instrumental in understanding some of the mechanisms by which resistance arises [34, 39], the genetic basis of resistance remains largely unresolved and is likely complex. For example, while resistance to benzimidazoles—the class of anthelmintics for which the genetic basis of resistance is best understood—has been linked clearly to mutations at three sites in the isotype-1 β-tubulin gene [40-42], there is evidence that it is a more complex trait than previously assumed [2]. In contrast, genome-wide studies of ivermectin response—another major anthelmintic—in a number of parasitic helminth species support the hypothesis that this is a quantitative, multigenic trait [43-45]. Therefore, establishing the genomic context in which drug resistance alleles are inherited using *H. contortus* will help to resolve the mechanisms by which resistance evolves and spreads in other species of parasitic nematodes as well.

The purpose of this study was to produce a genetic map of *H. contortus*, initially in order to establish an anchored framework for a draft genome under development, and subsequently to estimate the frequency and distribution of recombination in the genome. To do so, we performed a cross between two genetically divergent strains of *H. contortus* that differed in their anthelmintic resistance phenotypes: one that was fully susceptible and one that showed high levels of resistance to three commonly used anthelmintics [46, 47]. Four constraints restrict use of *H. contortus* crosses to implement standard classical approaches for genetic mapping: (i) there is an extremely high level of sequence polymorphism present both in field and laboratory strains of *H. contortus* [48] (ii) few very highly inbred isolates are available to use as parents, and so isolates comprise multiple genotypes; (iii) it is difficult, although not impossible, to perform single parent crosses from inbred lines [49, 50]; and (iv) mating is polyandrous, i.e. multiple males can and will mate with a single female [51]. We developed a genomic strategy for inferring segregation of single nucleotide polymorphisms within families by predicting paternal genotypes based on variants present in a single female and her progeny to construct an F_1_ genetic map. We discuss the implications of recombination, and other novel life history traits identified here, in the context of generating and maintaining genetic variation in parasite populations, and how these factors might impact the development and spread of anthelmintic resistance in this species.

## Results

### Genome sequencing and genetic diversity of a genetic cross between two isolates of *H. contortus*

A genetic cross was performed between two genetically and phenotypically defined *H. contortus* strains: females were from MHco3(ISE), a serially passaged anthelmintic susceptible “laboratory” strain that has been well characterised by genomic and transcriptomic analyses [24, 26], and males were from MHco18(UGA2004), a multi-drug resistant serially passaged strain originally isolated from the field at the University of Georgia, USA [46](Fig 1). Whole genome sequencing (WGS) was performed on DNA derived from a single adult MHco3(ISE) female parent and 41 of her F_1_ L_3_ progeny to achieve a minimum 30× sequencing coverage per sample (mean sequencing depth: 34.80× ± 16.16 standard deviations (SD)), generating a median yield of 65.97 million reads per sample (**S1 Table**). Mapping of the sequencing data was performed using an improved genome assembly of the MHco3(ISE) isolate described by Laing *et al.* [26], which now consists of five scaffolds representing the autosomal chromosomes and two scaffolds representing the X chromosome, for an assembly length of approximately 279 Mb. Sequence depth of the X chromosome scaffolds relative to the five autosomal scaffolds, together with rates of heterozygosity on the X chromosome scaffolds, revealed 20 male and 21 female F_1_ progenyin the brood.

**Fig 1.**
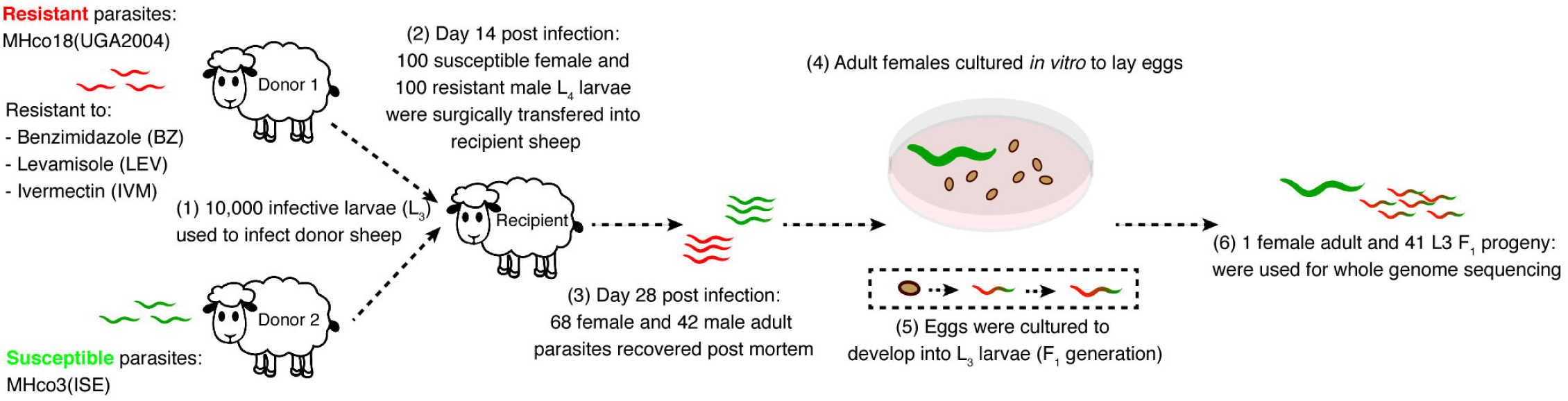
Outline of genetic cross between MHco3(ISE) drug susceptible and MHco18(UGA2004) multi-drug resistant *H. contortus*. A total of 68 MHco3(ISE) females and 42 MHco18(UGA2004) males (from an infection of 100 individuals of each sex) were recovered *post mortem*, after which reproductively mature females were incubated *in vitro* to lay eggs that were subsequently cultured to L_3_ stage. These larvae represent the F_1_ generation of the cross.

Approximately 5.3 million single nucleotide polymorphisms (SNPs) that passed stringent filtering criteria were identified in the autosomal chromosomes (Fig 2 A; **S2 Table**), at a genome-wide density of 2242 SNPs per 100 kb (Fig 2 B), or 1 SNP per 44.6 base pairs (bp). A pseudo-testcross approach was used to generate the F_1_ genetic map, which required that candidate markers: (i) were heterozygous in the female parent; and (ii) segregated in a ratio statistically indistinguishable from a 1:1 genotype ratio in the F_1_ progeny. By using these criteria, we identified a set of markers that could be analysed using the same statistical approaches as conventional linkage mapping using a test cross. Analysis of the 730,825 heterozygous SNPs in the female MHco3(ISE) parent demonstrated that the distribution of variation was not uniform throughout the genome, with a number of long contiguous regions of homozygosity observed (Fig 2 C; S1 Fig). In particular, approximately 27 Mb of the second half of chromosome IV was largely homozygous, containing about 50% more homozygous variant sites and about 30% less heterozygous sites compared to the genome-wide average (**S3 Table).**

**Fig 2.**
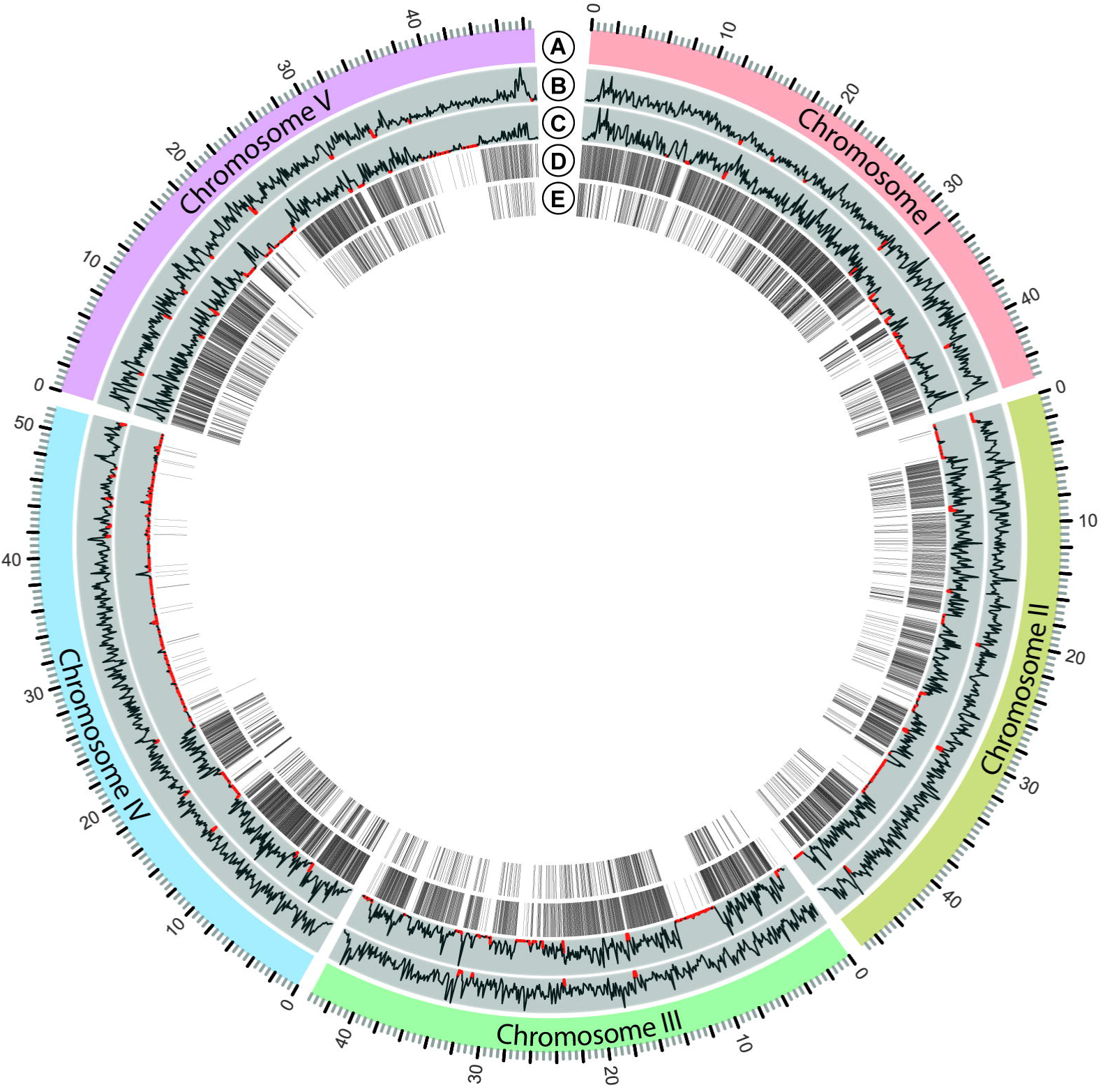
Autosome-wide variant density and candidate genetic map markers identified from the female parent and F_1_ progeny. (A) The five autosomes of *H. contortus*, named based on synteny with *C. elegans* chromosomes, span 237 Mb. (B) SNP density was calculated in 100 kbp windows, and is presented as the relative variant density of the female parent and all F_1_ progeny. (C) Density of heterozygous variants in the female parent. (D) Positions of candidate pseudo-testcross SNPs that were heterozygous in the female parent and segregated in a 1:1 genotype ratio in the F_1_ progeny. Red annotations in plots (C) and (D) highlight low density regions, defined as genome-wide mean SNP density minus 3 SD. (E) Positions of the final set of 1,618 SNPs used in the F_1_ genetic map. The plot was produced using *Circos* [52].

Among the SNPs that were heterozygous in the female parent, 171,876 SNPs segregated at an approximate 1:1 genotype ratio in the F_1_ progeny (**S2 Table**; PT:110 and PT:011). To avoid including tightly linked SNPs, the 171,876 candidate SNPs were thinned to 1 per 25,000 bp, which resulted in a final candidate list of 5,595 SNPs for analysis in the cross.

### Characterisation of an autosomal F_1_ genetic map generated using pseudo-testcross SNP markers

Initial analysis of genome-averaged genotype ratios (**S2 Fig**) of the candidate pseudo-testcross sites in each F_1_ individual revealed that most individuals displayed an approximate 50:50 ratio of homozygous:heterozygous genotypes, as expected. However, seven individuals presented as outliers with an excess of heterozygous genotypes (**S2 Fig A**; moderate outliers: individuals F1_12, F1_30, F1_40; extreme outliers: individuals F1_21, F1_23, F1_32, F1_38). The variant-allele frequency distribution of these individuals (**S3 Fig**) revealed a skew consistent with a non-diploid complement of chromosomes, with a major non-reference (relative to the genome assembly) allele frequency peak at approximately 30% and minor peak at 60% frequency. This allele frequency skew was typically found across all chromosomes within an individual, suggesting that they were not aneuploids. A notable exception was individual F1_30 (one of the moderate outliers), where chromosomes I, III, and V had a distinct allele frequency spectrum consistent with more than two copies of each chromosome present, relative to chromosomes II and IV, which appeared to be diploid. All seven of these non-diploid individuals were therefore removed from the pseudo-testcross analysis (**S2 Fig B, D; n = 34**).

A reanalysis of the remaining 34 individuals revealed 217,575 pseudo-testcross SNPs, 129,985 intercross SNPs, and 383,265 SNPS that were heterozygous in the female parent but did not segregate in a way compatible with analysis as a single-pair mating cross (**Table S2**). Thus, a total of 4,587 pseudo-testcross SNPs (217,575 SNPs thinned to 1 SNP per 25,000 bp) were candidate markers for the map construction using R/QTL (Fig 2 D), from which 1,618 SNPs were used in the final genetic map (Fig 2 E; Table S4). Recombination plots and genetic maps for the five autosomes are presented in Fig 3, and characteristics of the map are presented in Table 1. The total map distance of the five autosomes was approximately 344.46 cM. The number of markers per chromosome ranged from 215 on chromosome II to 475 on chromosome I, with a mean value of 323.6 markers per chromosome. Significant gaps in the map correlated with absence, or very low density, of the prerequisite heterozygous SNPs in the female parent, as described above (Fig 2 C). This loss of markers was most obvious in chromosome IV, where only approximately half of the chromosome is represented in the map, resulting in a map length of 49.21 cM, compared to the average map length of other chromosomes of 73.79 cM. The genome-wide recombination rate was on average 604.12 (± 84.01 SD) kb/cM or 1.68 (± 0.25 SD) cM/Mb, which corresponded to an overall average number of crossover events per chromosome of 0.69 (± 0.12 SD). Chromosome IV was again an outlier, with a recombination rate of 2.01 cM/Mb, approximately 21% higher than the other four autosomes (1.68 cM/Mb average).

**Table 1:** Summary characteristics of the F_1_ genetic map, including number of markers used, map length, recombination rate and crossover frequency.

| Chromosome | Chromosome length (bp) | Markers used (#) | Genetic map length (cM) | Recombination rate (Kb/cM) <sup>1</sup> | Recombination rate (cM/Mb) <sup>2</sup> | Crossovers per chromosome <sup>3</sup> |
| --- | --- | --- | --- | --- | --- | --- |
| I | 45778363 | 475 | 83.71 | 546.87 | 1.83 | 0.84 |
| II | 47384193 | 215 | 71.88 | 660.13 | 1.51 | 0.72 |
| III | 43564237 | 363 | 69.53 | 626.55 | 1.60 | 0.70 |
| IV <sup>4</sup> | 51819793 | 226 | 49.21 | 490.85 | 2.04 | 0.49 <sup>5</sup> |
| V | 48825595 | 339 | 70.13 | 696.22 | 1.44 | 0.70 |
| Total / average | 237372181 | 1618 | 344.46 | 604.12 | 1.68 | 0.69 |

1. Recombination rate (kb/cM): chromosome length (Kb) / genetic map length
2. Recombination rate (cM/Mb): genetic map length / (chromosomal length / 10<sup>6</sup>)
3. Crossovers per chromosome: ( genetic map length / 100 ) / number of chromosomes
4. The genetic map only spanned ~24 Mb of chromosome IV due to homozygosity in the female parent. As such, recombination rates have been calculated for chromosome IV using 24154752 bp (position of the genetic map marker closest to the homozygosity region) as the chromosome length.
5. Likely to be underestimated given only half of the chromosome is present.

**Fig 3.**
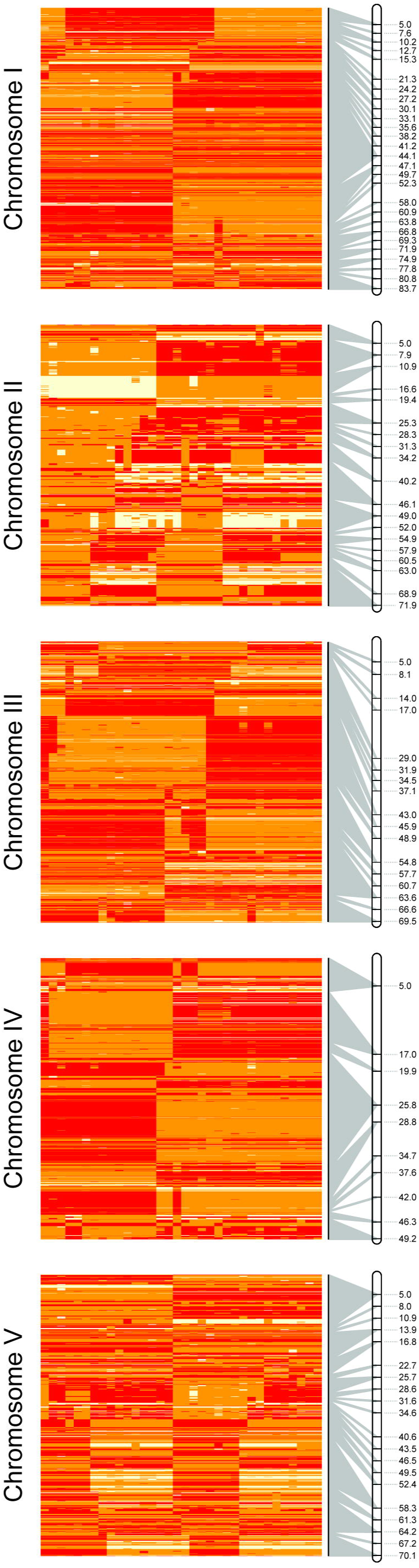
Recombination and genetic maps of the five autosomes of *H. contortus*. Recombination plots depict genotype segregation patterns per F_1_ progeny (columns; clustered by genetic similarity) of pseudo-testcross markers used in the genetic map (rows). Segregating “parental” and “recombinant” haplotypes inherited from the female parent are indicated by opposing colour schemes. Genotypes: AA: red; Aa: yellow; aa: white. The relationship between SNP position in the recombination map and genetic map position (cM) is represented by a connecting grey line; multiple SNPs between which no recombination was observed collapse into a single map position in the genetic map (grey ribbon from multiple SNPs to a single map marker).

Analysis of the X chromosome diversity from the adult female and all progeny revealed 100,016 SNPs in the 23.3 and 18.9 Mb X-linked scaffolds; this frequency (1 SNP per 422 bp) equates to approximately 10-fold fewer variable sites on the X chromosome relative to the autosomes. Attempts to generate an X chromosome genetic map were limited by a lack of prerequisite heterozygous variant sites in the female X chromosome sequences (**Fig S1**). To explore this further, the diversity of hemizygous genotypes called in the male F_1_ progeny, i.e. genotyped as AA or aa reflecting the haploid X^A^ or X^a^ allele, respectively, was compared to genotypes resolved in the female parent (**Fig S4**). Strikingly, male genotypes were entirely concordant with the female parent, further supporting the lack of segregating genetic diversity in the female parent diploid X chromosomes. Female F_1_ progeny contained both homozygous and heterozygous sites in their X chromosomes; given the lack of variation in the female parent, this diversity was entirely inherited from the paternal X chromosome.

### Patterns of recombination within autosomal chromosomes of the F_1_ progeny

Analysis of recombination rate throughout each chromosome was determined by comparing physical and genetic distances, which can be visualised in a Marey map [53](Fig. 4). Recombination rate (Fig. 4 red line; cM/Mb) was not uniform throughout the chromosomes, nor was it consistent between chromosomes. Chromosomes I, II and IV tended to show a pattern of three main recombination rate domains; a reduced recombination rate domain towards the middle of the chromosome, flanked by domains of increased recombination rate that extend toward the ends of the chromosomes. This three-domain pattern was not as clear for chromosomes III and V; chromosome III showed a greater recombination rate in the first half of the chromosome that decreased throughout the second half of the chromosome, whereas chromosome V had longer low recombination rate domains towards the ends of the chromosome arms, and greater recombination rate towards the middle of the chromosome. It is curious that chromosome IV retained the three-domain recombination architecture, given that the right arm is largely missing due to lack of the prerequisite heterozygous sites in this region of the female parent (Fig 2 C; S1 Fig). Each chromosome also showed evidence of additional low recombination rate domains at one or both ends of the chromosome in the sub-telomeric regions extending into the chromosome. Finally, within the elevated recombination rate domains, the recombination rate was not necessarily constant; discrete peaks of high recombination rates were observed in all chromosomes. However, the relative position of high recombination peaks was not the same between chromosomes.

**Fig 4.**
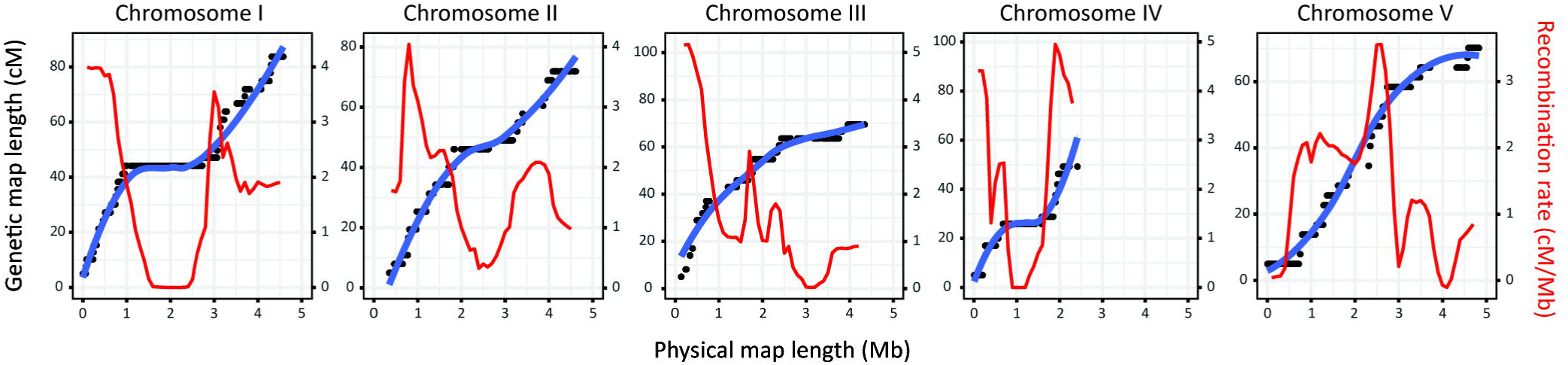
Analysis of recombination rate variation throughout the genome. Marey maps were constructed to show the relationship between the genetic position of each marker (black point) relative to the physical position of the marker in the genome. Line of best fit was plotted using default parameters of the *geom_smooth* function of *ggplot2* in R. Recombination rates (cM/Mb; red line) were calculated by calculating genetic map distance in 1 Mb windows throughout the genome from a fitted *loess*-smoothed line of the genetic map positions.

### Family structure and kinship among the brood

*H. contortus* is known to be polyandrous [51]. This knowledge, together with the observation that more than 50% of SNPs did not segregate in either a 1:1 or 1:2:1 genotype ratio (**Table S2**), suggested that the 41 progeny analysed were sired from more than a single male parent. An initial analysis of genetic relatedness by principal component analysis (PCA) of 21,822 autosomal SNPs (complete dataset thinned using a linkage disequilibrium threshold of 0.5 and minor allele frequency of 0.05) revealed obvious genetic structure, with at least four (PC 1 vs 2) to as many as six (PC 2 v 3) putative clusters of F_1_ progeny (Fig 5A), consistent with the hypothesis that the brood resulted from polyandrous mating.

**Fig 5.**
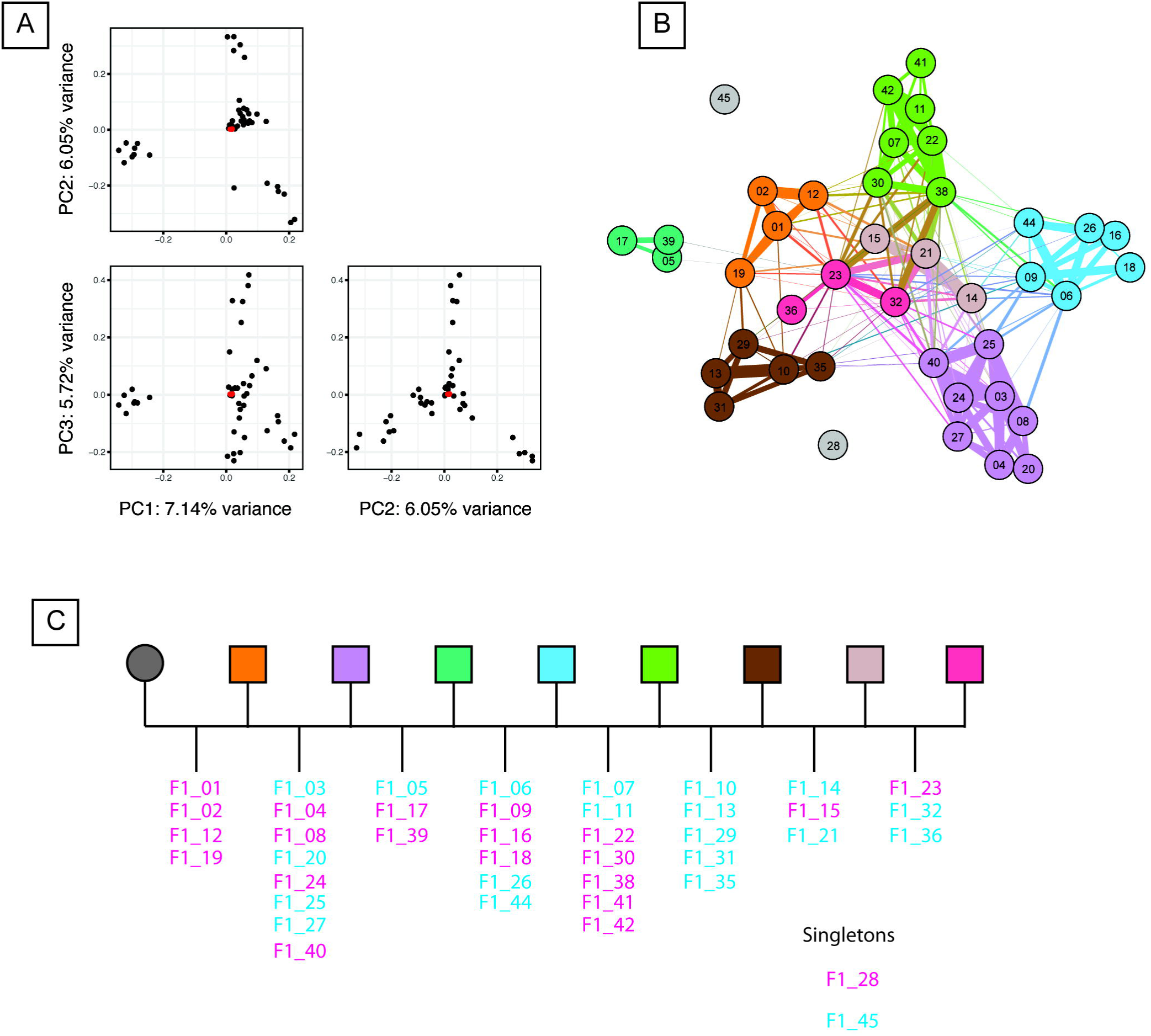
Familial relationships determined via analysis of genetic diversity and kinship between full- and half-sibs. (A) Principal component analysis of parent and progeny genetic diversity, comparing the top three principal components (PCA). The female parental values (n = 3) are indicated as red points in each plot. (B) Network analysis of kinship coefficients determined by *KING* [54] and visualised by *Gephi* [55] highlighting full-sib relationships between progeny. The thickness of the line (edges) represents the kinship coefficient between individuals (nodes) and is proportionate to the relationship between pairs. (C) Proposed pedigree of the brood. Full-sib male (blue) and female (pink) progeny are indicated for each sub-family. Colours used in (B) and (C) represent groups of progeny that share a common father.

To more accurately describe these putative relationships among the progeny, we calculated kinship coefficients [54], which describe the probability that a given allele in two individuals is identical by descent (i.e. an allele shared due to recent shared ancestry, as opposed to identical by state, in which the allele is simply shared by two individuals without common ancestry), for all pairwise combinations of progeny. Employing all autosomal SNPs (n = 5,323,039 SNPs), this analysis revealed eight clusters of full-sib relationships containing multiple F_1_ progeny (Fig 5B). Two individuals, F1_28 and F1_45, did not share any pairwise kinship coefficients consistent with a full-sib relationship with any individual, and hence, may represent the progeny from additional paternal contributions to the brood. Three individuals, F1_21, F1_23, and F1_38, seemed to show full-sib relationships with individuals from multiple families via strong kinship associations between themselves and others. Intriguingly, these were the same individuals identified as outliers with excess heterozygosity (**S2 Fig**) and that showed a skewed allele frequency distribution (**S3 Fig)** suggestive of aneuploidy or polyploidy. These autosomal kinship data are further supported by the observation that X chromosome diversity in the female F_1_ progeny, which reflects paternal X chromosome inheritance in the absence of maternal X chromosome diversity, clusters the female F_1_ progeny into five groups of two or more individuals (**S4 Fig**). Three unclustered individuals were also identified for the X chromosomes, including individual F1_28, which did not share any full-sib relationships in the kinship analysis (Fig 5B). These X chromosome derived clusters are concordant with the full-sib family structure using autosomal SNPs. Taken together, these data describing the familial relationships among the F_1_ progeny cohort lead us to propose a pedigree consisting of at least eight paternal contributions (Fig 5C).

## Discussion

Our comprehensive genetic characterisation of genome-wide patterns of segregation in progeny from a brood of parasites revealed extensive variation in recombination rates across chromosomes, and confirmed previous suggestions of polyandry as the dominant mating system in *H. contortus* [51]. Moreover, analysis of genetic variation in both autosomes and the X chromosome identified an extended region of reduced heterozygosity in the female parent, which could be a genetic consequence of population bottlenecks during the generation and maintenance of the MHco3(ISE) line. Analysis of allele frequency spectra also suggested the presence of polyploids among the progeny. The availability of a largely complete chromosomal scale *H. contortus* genome assembly facilitated such analyses. Here, we discuss some of the characteristics and challenges associated with the assembly of a genetic map when homozygous single parent crosses are not available, and how some of the features of the genetic cross impact on our understanding of *H. contortus* biology and anthelmintic resistance.

### Prediction of genomic structure

A small number of linkage maps have been described for free-living nematodes and parasitic helminths. *H. contortus* was found to have the lowest genome-wide recombination rate among these helminths, at an average of 604.12 kb/cM throughout the ∼280 Mb genome. However, the relative recombination rate (kb/cM) of *H. contortus* and other nematodes scales proportionately with genome size, i.e. larger genomes have lower recombination rates (**Fig S5**). While the recombination rates of some these nematodes are somewhat lower than predicted by a model describing the relationship between eukaryotic genome size and recombination rate (**Fig S5,** grey dashed line) [9], they are more consistent with recombination rates seen among other invertebrates (**Fig S5**, grey points; see Supplementary Table 1 from Lynch M [9] for invertebrate recombination rate data). The relationship between genome size and recombination rate is somewhat dependent on the number of crossovers per chromosome per meiosis; for example, in *C. elegans*, almost complete crossover interference occurs, such that only a single crossover per pair of homologous chromosome is observed [56]. In *H. contortus*, some but certainly not complete interference was observed, with an average rate of 0.69 crossovers per chromosome (i.e. 1.38 crossovers per pair of homologous chromosomes). This crossover rate is still substantially lower than in *S. mansoni*, whereby multiple chiasma per homologous pair have been observed [57], or in *M. hapla*, whereby recombination between all four chromatids within a homologous pair has been described [58]. The mechanisms by which this recombination rate diversity between helminth species is generated are largely unknown; however, it does provide an insight into the evolutionary potential of these diverse helminth species.

To our knowledge, we are the first to report the use of whole genome sequencing to construct a genetic map of any helminth species. WGS allowed significantly greater flexibility in choosing high quality variants to be included in the genetic map than other marker-based approaches such as amplified fragment length or Sanger-sequencing derived markers, and more recently, higher throughput RADseq and genotype-by-sequencing approaches, and allowed us to fully exploit the genetic variation in the available progeny. This was particularly important given that: (i) the progeny were not derived from a cross between genetically distinct homozygous single male and female parents, as is typical for a genetic mapping experiment; (ii) the high genetic diversity within isolates meant that a lot of markers have to be screened and discarded to find “bi-allelic markers” that segregate appropriately for analysis; and (iii) we did not know how many males would contribute to the progeny of the cross due to polyandry. As such, we developed a bioinformatics pipeline to select markers based on the genotype segregation ratio of the progeny (approximate 1:1 genotype ratios: Aa:aa [PT:011] or AA:Aa [PT:110]) and heterozygous sites in the female. This unusual cross design to account for the biological complexity meant that relatively few of the sites that differed between parents (pseudo-testcross SNPs represent only 4.09% of the total SNPs in the brood, and 29.77% of SNPs heterozygous in the female parent, before deliberate thinning) were usable in the map. A very large panel of traditional markers would thus have been required even for the relatively small number of progeny analysed here. The genome-wide resequencing approach that we used would seem to be the only practical way to generate complete recombination maps in this system. Genome-wide genetic variation that has been validated as segregating in a Mendelian fashion also provides a valuable resource for downstream experiments such as: QTL analyses of parasite traits (e.g. drug resistance); using individuals phenotyped *in vitro* using bioassays [59-62]; or as a source of genome-wide population genetic markers, which typically require low/no linkage disequilibrium between loci.

We initially intended to use the F_1_ genetic map to guide improvements of the assembly of the draft genome for *H. contortus* MHco3(ISE) [26]; while subsequent improvements to the genome assembly have rendered this unnecessary (unpublished data), the co-linearity of the genetic and physical maps confirms the accuracy of the current assembly. A number of features of this dataset would not have been obvious without integrating the genetic map and physical assembly. The first of these includes the non-uniform distribution of genetic map markers in the genome. This is most obvious in chromosome IV in which approximately half of the chromosome is missing from the genetic map, due to a long tract of homozygosity in the female parent. However, each chromosome contained multiple megabase-scale gaps that directly corresponded to a deficiency of heterozygosity in the female parent in these regions. This may reflect the genetic history of this particular strain: MHco3(ISE) is a laboratory strain that was originally generated by performing 15 rounds of half-sib matings of an outbred strain [47]; since that time, it has been passaged and cryopreserved on numerous occasions at an unknown, but likely limited, population size. Although significant diversity remains in this strain [63], it is probable that population bottlenecks, increased inbreeding or selection have resulted in discrete regions of the genome becoming genetically fixed. Secondly, the integration of the genetic map and contiguous physical genome map allowed us to describe the recombination landscape of the genome. Although there are similarities in the recombination rate domain structure with that of *C. elegans* [8, 14], chromosomes III and V have distinct recombination rate differences compared both to chromosomes I, II and IV of *H. contortus*, and to all chromosomes of *C. elegans*. The broad-scale distribution is unlikely to be the result of differential recombination around centromeric sequences, given the similarities in recombination domain structure with *C. elegans* chromosomes, and that *C. elegans* chromosomes are holocentric during mitosis [64, 65]. However, it has been proposed that the low or absent recombination in the chromosome termini may correlate with the presence of a spindle attachment site that guides segregation of homologous chromosomes in meiosis [66]. While we have no data to directly test whether *H. contortus* is holocentric, we have identified low recombining chromosome termini consistent with that observed in *C. elegans*.

Despite the relatively high marker density used here (n = 1,618), many SNPs were completely linked in seemingly non-recombining regions. Inclusion of a larger number of progeny would provide additional resolution to more precisely characterise variation in and transitions between recombination rate domains in each chromosome. Finally, although we could not generate a genetic map for the X chromosome due to the limited brood size and the absence of genetic diversity in the female parent, WGS data allowed us to examine genetic diversity among the female progeny, which highlighted both significant genetic variation and clustering consistent with shared paternal haplotypes in the autosomes.

### Detection of Polyandry

Technical challenges associated with single male and female mating led us to perform the genetic cross using 100 immature female MHco3(ISE) and 100 male MHco18(UGA2004) surgically implanted into the abomasum of a recipient sheep. Analysis of the genetic diversity among F_1_ progeny of a single female revealed discrete groups of progeny; given that *H. contortus* has been previously described to be polyandrous [51], we hypothesised that these groups represented the progeny of different male nematodes. In this cross, our data supports at least eight paternal genotypes contributing to multiple individuals in the brood (n = 41). These data are consistent with the original report of polyandry in *H. contortus*, which described at least 3 to 4 paternal microsatellite-derived genotypes from the 11 to 17 progeny sampled per single fecund female analysed [51]. Single worm genotyping of males recovered from the initial genetic cross recipient lamb would provide further insight into the ancestral relationships among the progeny. The relatively high frequency of polyandrous pairings would substantially increase the diversity of genotypes found among the progeny, as more possible pairs of haplotypes would be generated. This feature of *H. contortus* biology is likely to play a significant role in generating and maintaining the high levels of genetic diversity characterised in laboratory [63] and field [67, 68] isolates of this parasite and is also relevant to other parasitic nematode species where polyandry has been reported [69-71].

### Detection of non-diploid patterns of variation

*H. contortus* is a dioecious, sexually reproducing diploid animal. Unexpectedly, we observed seven of the 41 progeny (17.1%) with an excess of heterozygous genotypes, and with an allele frequency spectrum that is consistent with a polyploid complement of chromosomes. Moreover, two distinct patterns of allele frequency spectrum among six of the seven putative polyploids lead us to hypothesise that these progeny arose by either: (i) nondisjunction during meiosis 1 of gametogenesis in the female parent; or (ii) polyspermy, i.e. an egg that has been fertilized by more than one sperm, as a consequence of polyandry (see **Fig S6** for alternate hypotheses and evidence for the generation of triploid progeny in the brood). A third hypothesis—nondisjunction during male gametogenesis resulting in diploid sperm—was excluded; analysis of genotype frequencies among the F_1_ progeny at SNPs at which the female parent was homozygous demonstrated that paternally-derived alleles from putatively heterozygous sites were segregating independently, resulting in an approximate 1:1 genotype ratio among all but one individual (**Figure S2C;** the putative aneuploid F1_30). This supports the observation that polyploidy was inherited from diploid gametes derived from the female parent (i.e. nondisjunction), or multiple haploid gametes from the male parents (i.e. polyspermy).

Polyploidy has been previously described among nematodes. In *C. elegans*, a range of ploidy states have been characterised (see Hodgkin J [72] for review of work on natural and induced tetraploids, triploids and haploids) and is a feature of a cellular organismal growth into late adulthood due to nuclei endoreplication [73, 74]. However, polyploidy is typically associated with parthenogenesis in worms (e.g. some *Meloidogyne spp.* [75, 76] and some *Panagrolaimus spp.* [77]). Polyspermy in worms is thought to be rare, with a single description in the rodent filarial worm *Acanthocheilonema viteae* [78]; more is understood in regard to the mechanisms by which polyspermy is prevented [79-81]. However, polyspermy may be associated with polyandrous mating [82], whereby sexual conflict among males (at least 8 in the data presented) competing to reproduce with a female likely results in strong selection on male reproductive traits (e.g. sperm count, size and quality), which increases the likelihood of reproductive success [83]. While this would drive coevolution of female traits to block polyspermy, it may be that polyspermy is a consequence of this competition in polyandrous species such as *H. contortus*. Given that these progeny were sampled at the L_3_ stage, we cannot be sure that these individuals would have developed to adulthood and become reproductively viable. However, a report describing the karyotype of a single triploid *H. contortus* adult female suggests that they may be at least developmentally viable [23]. The presence of sporadic polyploidy among the *H. contortus* F_1_ progeny represents a novel finding among parasitic nematodes; further work is required to determine if triploidy is a feature of *H. contortus* biology and prevalent in the field, or, is a novel feature of this genetic cross. If the former is true, then it will be important to be aware of ploidy variation in population genetic studies of *H. contortus*, particularly if larval stages are sampled.

A single individual—F1_30—presented with a variant allele frequency spectrum consistent with an aneuploid complement of chromosomes. Aneuploidy and other severe chromosomal abnormalities have been described in experimental hybrid crosses between *H. contortus* and the related cattle parasite, *Haemonchus placei* [84]; such hybrids have recently been genetically characterized in the field [85]. Although such chromosomal abnormalities have not been described in within-species *H. contortus* crosses to date, the use of whole genome sequencing provides greater resolution over single marker techniques to detect these chromosome-wide changes, which may have resulted via incompatibility of rare alleles between the genetically diverse strains used in the cross.

## Conclusions

In summary, we have undertaken a comprehensive analysis of genetic diversity within a *H. contortus* family derived from an experimental genetic cross. Whole-genome sequencing of a female and her brood allowed the construction of a F_1_ genetic map, despite the challenging design dictated by the unusual biology and life history of this parasitic helminth. Development of the genetic map continues to build upon the genetic resources available for *H. contortus* as an experimentally tractable organism, and provides new insight into the recombination architecture of the genome. These data, together with evidence of polyandry and polyploidy, highlight the complexities of the underlying biology of *H. contortus*, and have important implications toward understanding the development and spread of anthelmintic resistance in this important pathogen of livestock. Clear recombination rate differences throughout the genome will influence the rate by which a locus correlated (i.e. a genetic marker linked to resistance), or causally associated (i.e. resistance conferring mutation) with anthelmintic resistance will evolve within a population, dependent on the position in the genome that the given locus lies. Incorporating recombination rate parameters in studies that aim to genetically detect or track the transmission of resistance will be critical to the utility and interpretation of data derived from such approaches. This will be particularly the case given the likely multigenic nature of resistance to some, and perhaps all, anthelmintics.

## Methods

### Construction of the genetic cross and collection of worm samples

A schematic of the experimental genetic cross is outlined in Fig 1. Briefly, two parasite naïve lambs were each infected with ∼10,000 infective larvae from one of two ovine-derived *H. contortus* strains, the anthelmintic susceptible MHco3(ISE) [47], or MHco18(UGA2004) [46], a multi-drug resistant strain that is insensitive to standard manufacturers recommended dose rates of benzimidazole, imidazothiazole and macrocyclic lactone anthelmintics. At 14 days post infection (DPI), developing sexually immature parasitic stages were recovered *post mortem*, and the sex of the L_4_ stage immature adults was determined by microscopic examination of gross morphology [86, 87]. A total of 100 MHco3(ISE) female and 100 MHco18(UGA2004) male L_4_ (F_0_ generation) were surgically transferred into the abomasum of a donor sheep to allow reproduction that would generate F_1_ hybrids between the two strains. At 28 DPI, 67 MHco3(ISE) females and 42 male MHco18(UGA2004) F_0_ from the recipient sheep were recovered *post mortem*, after which the males were snap frozen in liquid nitrogen and stored. Sampling was performed at 28 DPI to ensure that all of the females would have mated, and that they would be mature enough to have more viable progeny than is thought to be the case in early patency. Individual females were placed into individual wells of 24-well cluster plates (Sarstedt) containing 1 mL of warm RPMI 1640 cell culture media containing 1% (v/v) D-glucose, 2 mM glutamine, 100 IU/mL penicillin, 100 mg/mL streptomycin, 125 mg/mL gentamycin, 25 mg/mL amphotericin B [88] and Hepes (1% v/v) and incubated in 5% CO_2_ at 37° C for 48 h to promote egg shedding. Eggs were transferred at 24 h and 48 h and mixed with fresh helminth egg-free sheep faeces before being incubated at 24° C for 2 weeks to allow larval development to L_3_. After this time, a single female parent (F_0_) and a total of 41 F_1_ L_3_ progeny were individually stored in preparation for DNA extraction and sequencing library preparation.

### Sample preparation and sequencing

The female parent was dissected on ice to isolate the head and anterior body only (in three sections, as three technical replicates) to avoid contamination with fertilised eggs present *in utero*. The female sections and individual L_3_ were transferred into 10 μL of sample lysis buffer (working solution: 1000 μL Direct PCR Lysis Reagent [Viagen, Los Angeles, USA], 50 μL 1 M DTT, 10 μL 100 mg/ml Proteinase K) in a 96-well plate and allowed to incubate at 60°C for 2 h followed by 85°C for 45 min. Whole genome amplification (WGA) of each sample lysate was performed using RepliG amplification. First, 2-5 μL of sample lysate was combined with 5 μL of 1.3 M Trehalose in a 96-well plate and mixed by gentle tapping, incubated for 3 min at 95°C, and placed on ice. A 40 μL RepliG amplification mix (29 μL REPLI-g Reaction Buffer + 1 μL REPLI-g polymerase + 10 μL 1.3 M Trehelose) was added to each well, and incubated for 16 h at 30°C followed by 10 min at 65°C before being placed on ice. The WGA DNA was cleaned using Ampure XP beads at a 1.4× bead:DNA reaction ratio, before being eluted in 50 μL of RNase/DNase-free water and stored at 4°C.

PCR-free sequencing libraries (mean length of approximately 400 bp) were prepared by methods previously described [89] and sequenced on an Illumina HiSeq X10, resulting in approximately 3.06×10^9^ 151-bp paired-end reads (see **S1 Table** for a breakdown of reads per lane and per sample). Raw sequence data is archived under the ENA study accession ERP024253.

### Mapping and variant analysis

Raw sequence data was mapped to the current unpublished version of the reference genome for *Haemonchus contortus* (v3.0, available at ftp://ngs.sanger.ac.uk/production/pathogens/Haemonchus_contortus) using *Smalt* (http://www.sanger.ac.uk/science/tools/smalt-0) with the mapping parameters “-y 0.8 -i 800”. Data from multiple sequencing lanes for a single sample were merged (*samtools-1.3 merge*) and duplicate reads removed (*Picard v2.5.0*; https://github.com/broadinstitute/picard) from the bam files before further processing. Variants were called using *GATK Unified Genotyper* (v3.3.0)[90]. The raw variant set was initially filtered to flag variants as low quality if they met the following conditions: quality by depth (QD) < 2; Fisher’s test of strand bias (FS) > 60; RMS mapping quality (MQ) < 40; rank sum of alt vs reference mapping quality (MQRankSum) < −12.5; read position rank sum (ReadPosRankSum) < 8; read depth (DP) < 10. Variants were filtered further using *vcftools* (v0.1.14)[91] to exclude sites with low quality flags, minimise loci with missing data *(“--max missing 0.8”*), exclude indels (“*--remove-indels”*), exclude SNPs with genotype quality (GQ) < 30, and ensure sites were biallelic *(“--min-alleles 2, --max-alleles 2”*). A gff file generated from *RepeatMasker* of the reference genome was also used to filter variants from the vcf file that were likely associated with repetitive and difficult to map regions.

Sex determination of the F_1_ progeny was performed by measuring: (i) the relative autosome to X chromosome (characterised and thus named based on synteny with *C. elegans* autosomes and X chromosome) read depth using *samtools-1.3 bedcov*; and (ii) the relative heterozygosity of the X chromosome using *vcftools* (v0.1.14) “*--het”*.

### Genetic map construction

A “pseudo-testcross” (PT) strategy [92] was employed to generate the genetic map, which required that each input variant site was: (i) heterozygous in the female parent, and (ii) segregating in a 1:1 genotype ratio in the F_1_ progeny. The segregation pattern of each SNP was first calculated in the F_1_ progeny (with “A: referring to the reference allele and “a” to the alternative), which resulted in SNPs being placed into one of four categories that best described the likely genotypes of the parents of the cross for that given SNP: (i) “PT:110”, i.e. AA×Aa, (ii) “PT:011”, i.e. Aa×aa, (iii) “intercross”, i.e. Aa×Aa, or (iv) SNPs that were clearly segregating in the brood, but for which the segregation ratio of genotypes in the progeny did not fit a simple Mendelian segregation pattern that could be generated via reproduction from a single pair of parents. SNP density was further reduced using *vcftools* (v0.1.14)[91] *--thin* as described in the text. The number of filtered SNPs per segregation group is described in **S2 Table.** Genotypes for autosomal PT:011 and PT:110 SNPs that were heterozygous in the female parent were imported into *R-3.2.2* [93], after which pairwise recombination fractions (RF) and logarithm of the odds (LOD) scores were determined for each chromosome using *R/QTL* [94]. Recombination fractions were converted into map distance in centimorgans (cM) using the kosambi map function. Variants resulting in inflation of map distances were identified using *qtl::droponemarker*, and as outliers relative to surrounding markers via visual inspection of LOD and RF using *qtlcharts*::*iPlot* [95]. These aberrant markers were removed in the generation of the final map.

A reverse cross design, whereby SNPs were chosen that: (i) segregated in a 1:1 genotype ratio; and (ii) were homozygous in the female parent (and therefore putatively heterozygous in the male parents) was also performed. Although polyandry prevented a male-specific genetic map from being constructed (multiple male parents confounded the calculation of linkage between heterozygous sites), these data were used to determine the segregation frequency of alleles from the male parents.

### Recombination landscape

Recombination patterns for each chromosome were visualised first by generating genotype matrices of pseudo-testcross markers for each chromosome using *vcftools* (v0.1.14) “*--012”*, followed by plotting using the *gplots:*:*heatmap2* function in R. These maps highlighted recombination breakpoints, linkage blocks, and regions of excess heterozygosity or reduced heterozygosity. Recombination rate changes throughout the genome were visualised by constructing Marey maps, which compare the position of the marker in the genome (base position in the fasta sequence) to the relative position in the genetic map. A fitted loess smoothed line of the genetic map positions in 1 Mb windows was performed to calculate the recombination rate.

### Kinship analysis

Analysis of genetic relatedness between F_1_ progeny was undertaken to characterise evidence of polyandry and to determine, if present, the impact on the cross analysis. Principal component analysis (PCA) of genetic distances between the F_1_ progeny and female parent was performed using the *SNPrelate* package in R 3.1.2 [96]. Kinship coefficients were determined for all pairwise relationships among the F_1_ progeny using *KING* [54]. Relationship networks of the pairwise kinship coefficients were visualised using *Gephi* (v 0.9.1; [55]) to highlight full- and half-sib relationships among the F_1_ progeny. Layout of the kinship network graph was determined using the *Force Atlas* parameter, with the nodes (Findividuals) coloured by their proposed kinship group, and the thickness of the edges proportionate to the kinship coefficient between two F_1_ individuals (nodes).

## List of abbreviations

cM: centimorgan
DPI: days post infection
LOD: logarithm of the odds
MHco3(ISE): inbred susceptible *H. contortus* strain
MHco18(UGA2004): triple anthelmintic resistant *H. contortus* strain
PCA: principle components analysis
PT: pseudo-testcross
QTL: quantitative trait loci
RF: recombination fraction
SNP: single nucleotide polymorphism
WGA: whole genome amplification
WGS: whole genome sequencing.

## Declarations

### Ethics approval and consent to participate

All experimental procedures described in this manuscript were examined and approved by the Moredun Research Institute Experiments and Ethics Committee and were conducted under approved UK Home Office licenses in accordance with the Animals (Scientific Procedures) Act of 1986. The Home Office licence number is PPL 60/03899 and experimental code identifier was E46/11.

### Consent for publication

Not applicable

### Availability of data and material

The raw sequencing data generated and/or analysed during the current study are available in the European Nucleotide Archive repository, http://www.ebi.ac.uk/ena/ under the study accession number ERP024253. The genome assembly is available at ftp://ngs.sanger.ac.uk/production/pathogens/Haemonchus_contortus.

### Author’s contributions

Conceived the study: ED, RL, AT, JAC, JSG, NDS

Undertook the genetic cross: NDS, DJB, AAM

Performed the molecular biology: KM, RL

Coordinated sequencing: NH

Participated in the discussion and interpretation of results: SRD, RL, DJB, CB, UC, JSG, NH,

BKM, KM, AAM, AT, AT, MB, ED, JAC, NDS

Performed the data analysis: SRD

Wrote the first draft of the manuscript: SRD, JAC

All authors read and critically revised the final manuscript.

## Acknowledgements

We would like to acknowledge the BUG Consortium members and Parasite Genomics group (WTSI) for helpful comments and suggestions toward this work, Pathogen Informatics and DNA Pipelines (WTSI) for their support and expertise, Taisei Kikuchi for the whole genome amplification protocol for nematode larvae, Ray Kaplan for supplying L_3_ larvae from the MHco18(UGA2004) strain, and to the Bioservices Division, Moredun Research Institute, for expert care and assistance with animals.

## Supporting information

**S1 Fig. Genome-wide variant density of the female parent.** SNP density is presented as the number of homozygous reference (AA; panel A), heterozygous (Aa; panel B) and homozygous variant (aa; panel C) SNPs per 100-kbp. Plots are coloured per chromosome, in the following order: I (black), II (red), III (green), IV (dark blue), and V (light blue). The X chromosome has also been included (currently in two scaffolds) as indicated by the purple and yellow segments.

**S2 Fig. Genome-wide average pseudo-testcross SNP density in the F_1_ progeny.** Pseudo-testcross markers were chosen based on an approximate segregation ratio of 1:1 homozygous:heterozygous genotypes among the F_1_ progeny. Analysis of heterozygous Aa genotype frequencies (A) of the 41 F_1_ progeny revealed a number of individuals presenting with moderate and extreme heterozygosity. A reanalysis of Aa genotype frequencies after the outlier individuals were removed (34 individuals remaining) (B) resulted in genotype frequencies at approximate 1:1 genotype ratio. (C) Comparison of heterozygosity among the F1 progeny at SNPs selected that segregate at a 1:1 genotype ratio in the progeny and are homozygous in the female parent; these sites are therefore putatively heterozygous in the male parents; i.e. a reverse F_1_ cross. In this comparison, only a single F_1_ individual – F1_30 – showed moderate heterozygosity. Individual points are coloured based on deviation from null expectation (H_0_: 1:1 genotype ratio of homozygous:heterozygous sites) determined by chi-square analysis (*X*^2^, df=1). Median frequency (solid grey line) and “whiskers” (dashed grey lines; most extreme point no more than 1.5× the interquartile range) were calculated using the R function *boxplot.stats*.

**S3 Fig. Variant allele frequency density plots used to explore ploidy among the F_1_ progeny.** Each plot displays the variant allele frequency of each chromosome (coloured lines) and genome-wide average (black dotted line).

**S4 Fig. X chromosome genetic diversity of 41 F^1^ progeny and female parent (3 replicate samples).** SNPs genotyped as hemizygous in male samples were analysed in all samples to detect segregation of X chromosome variants from the female parent. Female genotypes: X^A^ X^A^: red; X^A^ X^a^: yellow; X^a^ X^a^: white. Male genotype (hemizygous): X^A^ O: red; X^a^ O: white.

**S5 Fig. Relationship between recombination rate (kb/cM) and genome size (Mb)**. Recombination rates for helminth species with published genetic maps—*Caenorhabditis elegans* [14], *Haemonchus contortus* (current study), *Meloidogyne hapla* [5, 6], Pristionchus pacificus [4], and *Schistosoma mansoni* [17] —were derived from known genome size and the reported genetic map length. These estimates were compared against a derivation of the equation presented by Lynch M [9] describing the relationship between recombination rate and genome size (recombination rate (cM/Mb) = 0.0019×[genome_size(Mb)] ^-0.71^. These data were converted to kb/cM (1/[cM/Mb]×1000). Based on this equation, recombination rates were estimated for genome sizes between 10 and 1000 Mb (grey dashed line). The helminth and modelled data were plotted with recombination rates and genome sizes of invertebrate species presented in Supplementary Table 1of Lynch M [9].

**S6 Fig. Alternate hypotheses proposed to explain the segregation of alleles and recombination, and the presence of triploid progeny.** Four hypotheses for the segregation of genetic variation in gametes produced from the heterozygous female are presented: (1) normal gametogenesis; (2) nondisjunction in meiosis 1; (3) nondisjunction in meiosis 2; and (4) polyspermy.

**S1 Table. Sequencing data used in this study.**

**S2 Table. Breakdown of genetic variation in the female parent, and proportion of variants in each segregation class.**

**S3 Table. Genotype concordance between three female parent (replicate) samples.**

**S4 Table. SNPs used in the final genetic map.**

**S5 Table. Expected and observed genetic consequences of triploidy via nondisjunction or polyspermy in the cross.** Three alternate hypotheses and data are presented: (1) expected segregation of pseudo-testcross markers, which were used in the making of the genetic map; (2) triploidy via nondisjunction; and (3) triploidy via polyspermy.

## References

1. Valentim CL, Cioli D, Chevalier FD, Cao X, Taylor AB, Holloway SP, Pica-Mattoccia L, Guidi A, Basso A, Tsai IJ, et al: Genetic and molecular basis of drug resistance and species-specific drug action in schistosome parasites. Science 2013, 342:1385–1389.

2. Zamanian M, Cook DE, Lee D, Lee J, Andersen E: Heritable Small RNAs Regulate Nematode Benzimidazole Resistance. bioRxiv 2017.

3. Srinivasan J, Sinz W, Jesse T, Wiggers-Perebolte L, Jansen K, Buntjer J, van der Meulen M, Sommer RJ: An integrated physical and genetic map of the nematode Pristionchus pacificus. Mol Genet Genomics 2003, 269:715–722.

4. Srinivasan J, Sinz W, Lanz C, Brand A, Nandakumar R, Raddatz G, Witte H, Keller H, Kipping I, Pires-daSilva A, et al: A bacterial artificial chromosome-based genetic linkage map of the nematode Pristionchus pacificus. Genetics 2002, 162:129–134.

5. Thomas VP, Fudali SL, Schaff JE, Liu Q, Scholl EH, Opperman CH, Bird DM, Williamson VM: A sequence-anchored linkage map of the plant-parasitic nematode Meloidogyne hapla reveals exceptionally high genome-wide recombination. G3 (Bethesda) 2012, 2:815–824.

6. Opperman CH, Bird DM, Williamson VM, Rokhsar DS, Burke M, Cohn J, Cromer J, Diener S, Gajan J, Graham S, et al: Sequence and genetic map of Meloidogyne hapla: A compact nematode genome for plant parasitism. Proc Natl Acad Sci U S A 2008, 105:14802–14807.

7. Nemetschke L, Eberhardt AG, Viney ME, Streit A: A genetic map of the animal-parasitic nematode Strongyloides ratti. Mol Biochem Parasitol 2010, 169:124–127.

8. Rockman MV, Kruglyak L: Recombinational landscape and population genomics of Caenorhabditis elegans. PLoS Genet 2009, 5:e1000419.

9. Lynch M: The origins of eukaryotic gene structure. Mol Biol Evol 2006, 23:450–468.

10. Burt A, Bell G, Harvey PH: Sex differences in recombination. J Evol Biol 1991, 4:259–277.

11. Lenormand T, Dutheil J: Recombination difference between sexes: a role for haploid selection. PLoS Biol 2005, 3:e63.

12. Beye M, Gattermeier I, Hasselmann M, Gempe T, Schioett M, Baines JF, Schlipalius D, Mougel F, Emore C, Rueppell O, et al: Exceptionally high levels of recombination across the honey bee genome. Genome Res 2006, 16:1339–1344.

13. Tortereau F, Servin B, Frantz L, Megens HJ, Milan D, Rohrer G, Wiedmann R, Beever J, Archibald AL, Schook LB, Groenen MAM: A high density recombination map of the pig reveals a correlation between sex-specific recombination and GC content. BMC Genomics 2012, 13.

14. Barnes TM, Kohara Y, Coulson A, Hekimi S: Meiotic recombination, noncoding DNA and genomic organization in Caenorhabditis elegans. Genetics 1995, 141:159–179.

15. Chan AH, Jenkins PA, Song YS: Genome-Wide Fine-Scale Recombination Rate Variation in Drosophila melanogaster. PLoS Genet 2012, 8.

16. Kaur T, Rockman MV: Crossover heterogeneity in the absence of hotspots in Caenorhabditis elegans. Genetics 2014, 196:137–148.

17. Criscione CD, Valentim CL, Hirai H, LoVerde PT, Anderson TJ: Genomic linkage map of the human blood fluke Schistosoma mansoni. Genome Biol 2009, 10:R71.

18. Cutter AD, Payseur BA: Selection at linked sites in the partial selfer Caenorhabditis elegans. Mol Biol Evol 2003, 20:665–673.

19. Rockman MV, Skrovanek SS, Kruglyak L: Selection at linked sites shapes heritable phenotypic variation in C. elegans. Science 2010, 330:372–376.

20. Andersen EC, Gerke JP, Shapiro JA, Crissman JR, Ghosh R, Bloom JS, Felix M-A, Kruglyak L: Chromosome-scale selective sweeps shape Caenorhabditis elegans genomic diversity. Nat Genet 2012, 44:285–290.

21. Urquhart GM: Veterinary parasitology. 2nd edn. Oxford, UK; Ames, Iowa: Blackwell Science; 1996.

22. Gilleard JS: Haemonchus contortus as a paradigm and model to study anthelmintic drug resistance. Parasitology 2013, 140:1506–1522.

23. Bremner KC: Cytological polymorphism in the nematode Haemonchus contortus (Rudolphi 1803) Cobb 1898. Nature 1954, 174:704–705.

24. Laing R, Martinelli A, Tracey A, Holroyd N, Gilleard JS, Cotton JA: Haemonchus contortus: Genome Structure, Organization and Comparative Genomics. Adv Parasitol 2016, 93:569–598.

25. Schwarz EM, Korhonen PK, Campbell BE, Young ND, Jex AR, Jabbar A, Hall RS, Mondal A, Howe AC, Pell J, et al: The genome and developmental transcriptome of the strongylid nematode Haemonchus contortus. Genome Biol 2013, 14:R89.

26. Laing R, Kikuchi T, Martinelli A, Tsai IJ, Beech RN, Redman E, Holroyd N, Bartley DJ, Beasley H, Britton C, et al: The genome and transcriptome of Haemonchus contortus, a key model parasite for drug and vaccine discovery. Genome Biol 2013, 14:R88.

27. Le Jambre LF, Gill JH, Lenane IJ, Baker P: Inheritance of avermectin resistance in Haemonchus contortus. Int J Parasitol 2000, 30:105–111.

28. Le Jambre LF: Genetics of vulvar morph types in Haemonchus contortus: Haemonchus contortus cayugensis from the Finger Lakes Region of New York. Int J Parasitol 1977, 7:9–14.

29. Le Jambre LF, Royal WM, Martin PJ: The inheritance of thiabendazole resistance in Haemonchus contortus. Parasitology 1979, 78:107–119.

30. Sangster NC, Redwin JM, Bjorn H: Inheritance of levamisole and benzimidazole resistance in an isolate of Haemonchus contortus. Int J Parasitol 1998, 28:503–510.

31. Hunt PW, Kotze AC, Knox MR, Anderson LJ, McNally J, Lej LF: The use of DNA markers to map anthelmintic resistance loci in an intraspecific cross of Haemonchus contortus. Parasitology 2010, 137:705–717.

32. Le Jambre LF, Geoghegan J, Lyndal-Murphy M: Characterization of moxidectin resistant Trichostrongylus colubriformis and Haemonchus contortus. Vet Parasitol 2005, 128:83–90.

33. Redman E, Sargison N, Whitelaw F, Jackson F, Morrison A, Bartley DJ, Gilleard JS: Introgression of ivermectin resistance genes into a susceptible Haemonchus contortus strain by multiple backcrossing. PLoS Pathog 2012, 8:e1002534.

34. Kotze AC, Hunt PW, Skuce P, von Samson-Himmelstjerna G, Martin RJ, Sager H, Krucken J, Hodgkinson J, Lespine A, Jex AR, et al: Recent advances in candidate-gene and whole-genome approaches to the discovery of anthelmintic resistance markers and the description of drug/receptor interactions. Int J Parasitol Drugs Drug Resist 2014, 4:164–184.

35. Mederos AE, Ramos Z, Banchero GE: First report of monepantel Haemonchus contortus resistance on sheep farms in Uruguay. Parasit Vectors 2014, 7:598.

36. Sales N, Love S: Resistance of Haemonchus sp. to monepantel and reduced efficacy of a derquantel / abamectin combination confirmed in sheep in NSW, Australia. Vet Parasitol 2016, 228:193–196.

37. Kaminsky R, Ducray P, Jung M, Clover R, Rufener L, Bouvier J, Weber SS, Wenger A, Wieland-Berghausen S, Goebel T, et al: A new class of anthelmintics effective against drug-resistant nematodes. Nature 2008, 452:176–180.

38. Bassetto CC, Amarante AF: Vaccination of sheep and cattle against haemonchosis. J Helminthol 2015, 89:517–525.

39. Britton C, Roberts B, Marks ND: Functional Genomics Tools for Haemonchus contortus and Lessons From Other Helminths. Adv Parasitol 2016, 93:599–623.

40. Kwa MS, Veenstra JG, Roos MH: Benzimidazole resistance in Haemonchus contortus is correlated with a conserved mutation at amino acid 200 in beta-tubulin isotype 1. Mol Biochem Parasitol 1994, 63:299–303.

41. Silvestre A, Cabaret J: Mutation in position 167 of isotype 1 beta-tubulin gene of Trichostrongylid nematodes: role in benzimidazole resistance? Mol Biochem Parasitol 2002, 120:297–300.

42. Ghisi M, Kaminsky R, Maser P: Phenotyping and genotyping of Haemonchus contortus isolates reveals a new putative candidate mutation for benzimidazole resistance in nematodes. Vet Parasitol 2007, 144:313–320.

43. Doyle SR, Bourguinat C, Nana-Djeunga HC, Kengne-Ouafo JA, Pion SDS, Bopda J, Kamgno J, Wanji S, Che H, Kuesel AC, et al: Genome-wide analysis of ivermectin response by Onchocerca volvulus reveals that genetic drift and soft selective sweeps contribute to loss of drug sensitivity. PLoS Negl Trop Dis 2017.

44. Bourguinat C, Lee AC, Lizundia R, Blagburn BL, Liotta JL, Kraus MS, Keller K, Epe C, Letourneau L, Kleinman CL, et al: Macrocyclic lactone resistance in Dirofilaria immitis: Failure of heartworm preventives and investigation of genetic markers for resistance. Vet Parasitol 2015, 210:167–178.

45. Choi YJ, Bisset SA, Doyle SR, Hallsworth-Pepin K, Martin J, Grant WN, Mitreva M: Genomic introgression mapping of field-derived multiple-anthelmintic resistance in the nematode parasite Teladorsagia circumcincta. PLoS Genet 2017, 13:e1006857.

46. Williamson SM, Storey B, Howell S, Harper KM, Kaplan RM, Wolstenholme AJ: Candidate anthelmintic resistance-associated gene expression and sequence polymorphisms in a triple-resistant field isolate of Haemonchus contortus. Mol Biochem Parasitol 2011, 180:99–105.

47. Roos MH, Otsen M, Hoekstra R, Veenstra JG, Lenstra JA: Genetic analysis of inbreeding of two strains of the parasitic nematode Haemonchus contortus. Int J Parasitol 2004, 34:109–115.

48. Gilleard JS, Redman E: Genetic Diversity and Population Structure of Haemonchus contortus. Adv Parasitol 2016, 93:31–68.

49. Sargison N, Redman E, Morrison AA, Bartley DJ, Jackson F, Naghra-van Gijzel H, Holroyd N, Berriman M, Cotton JA, Gilleard JS: Single pair mating as a method to reduce genetic polymorphism in the parasitic nematode Haemonchus contortus. submitted to XXX; personal communication 2017.

50. Sargison N: Development of genetic crossing methods to identify genes associated with macrocyclic lactone resistance in the sheep nematode parasite, Haemonchus contortus. University of Edinburgh, 2009.

51. Redman E, Grillo V, Saunders G, Packard E, Jackson F, Berriman M, Gilleard JS: Genetics of mating and sex determination in the parasitic nematode Haemonchus contortus. Genetics 2008, 180:1877–1887.

52. Krzywinski MI, Schein JE, Birol I, Connors J, Gascoyne R, Horsman D, Jones SJ, Marra MA: Circos: An information aesthetic for comparative genomics. Genome Res 2009, 19:1639–1645.

53. Chakravarti A: A graphical representation of genetic and physical maps: the Marey map. Genomics 1991, 11:219–222.

54. Manichaikul A, Mychaleckyj JC, Rich SS, Daly K, Sale M, Chen WM: Robust relationship inference in genome-wide association studies. Bioinformatics 2010, 26:2867–2873.

55. Bastian M, Heymann S, Jacomy M: Gephi: an open source software for exploring and manipulating networks. ICWSM 2009, 8:361–362.

56. Meneely PM, Farago AF, Kauffman TM: Crossover distribution and high interference for both the X chromosome and an autosome during oogenesis and spermatogenesis in Caenorhabditis elegans. Genetics 2002, 162:1169–1177.

57. Hirai H, Hirata M, Aoki Y, Tanaka M, Imai HT: Chiasma analyses of the parasite flukes, Schistosoma and Paragonimus (Trematoda), by using the chiasma distribution graph. Genes Genet Syst 1996, 71:181–188.

58. Liu QLL, Thomas VP, Williamson VM: Meiotic parthenogenesis in a root-knot nematode results in rapid genomic homozygosity. Genetics 2007, 176:1483–1490.

59. Hunt KR, Taylor MA: Use of the egg hatch assay on sheep faecal samples for the detection of benzimidazole resistant nematodes. Vet Rec 1989, 125:153–154.

60. Le Jambre LF: Egg hatch as an in vitro assay of thiabendazole resistance in nematodes. Vet Parasitol 1976, 2:385–391.

61. Coles GC, Tritschler JP, 2nd, Giordano DJ, Laste NJ, Schmidt AL: Larval development test for detection of anthelmintic resistant nematodes. Res Vet Sci 1988, 45:50–53.

62. Alvarez-Sanchez MA, Perez Garcia J, Bartley D, Jackson F, Rojo-Vazquez FA: The larval feeding inhibition assay for the diagnosis of nematode anthelmintic resistance. Exp Parasitol 2005, 110:56–61.

63. Redman E, Packard E, Grillo V, Smith J, Jackson F, Gilleard JS: Microsatellite analysis reveals marked genetic differentiation between Haemonchus contortus laboratory isolates and provides a rapid system of genetic fingerprinting. Int J Parasitol 2008, 38:111–122.

64. Wicky C, Rose AM: The role of chromosome ends during meiosis in Caenorhabditis elegans. Bioessays 1996, 18:447–452.

65. Albertson DG, Thomson JN: Segregation of holocentric chromosomes at meiosis in the nematode, Caenorhabditis elegans. Chromosome Res 1993, 1:15–26.

66. Cutter AD, Dey A, Murray RL: Evolution of the Caenorhabditis elegans genome. Mol Biol Evol 2009, 26:1199–1234.

67. Redman E, Whitelaw F, Tait A, Burgess C, Bartley Y, Skuce PJ, Jackson F, Gilleard JS: The Emergence of Resistance to the Benzimidazole Anthlemintics in Parasitic Nematodes of Livestock Is Characterised by Multiple Independent Hard and Soft Selective Sweeps. PLoS Negl Trop Dis 2015, 9:e0003494.

68. Silvestre A, Sauve C, Cortet J, Cabaret J: Contrasting genetic structures of two parasitic nematodes, determined on the basis of neutral microsatellite markers and selected anthelmintic resistance markers. Mol Ecol 2009, 18:5086–5100.

69. Zhou CH, Yuan K, Tang XL, Hu NY, Peng WD: Molecular genetic evidence for polyandry in Ascaris suum. Parasitol Res 2011, 108:703–708.

70. Hildebrandt JC, Eisenbarth A, Renz A, Streit A: Single worm genotyping demonstrates that Onchocerca ochengi females simultaneously produce progeny sired by different males. Parasitol Res 2012, 111:2217–2221.

71. Grillo V: Development of microsatellites and the population genetic analysis of the parasitic nematode Teladorsagia circumcincta. University of Glasgow, 2006.

72. Hodgkin J: Karyotype, ploidy, and gene dosage. WormBook 2005:1–9.

73. Flemming AJ, Shen ZZ, Cunha A, Emmons SW, Leroi AM: Somatic polyploidization and cellular proliferation drive body size evolution in nematodes. Proc Natl Acad Sci U S A 2000, 97:5285–5290.

74. Hedgecock EM, White JG: Polyploid tissues in the nematode Caenorhabditis elegans. Dev Biol 1985, 107:128–133.

75. Lunt DH, Kumar S, Koutsovoulos G, Blaxter ML: The complex hybrid origins of the root knot nematodes revealed through comparative genomics. PeerJ 2014, 2:e356.

76. Blanc-Mathieu R, Perfus-Barbeoch L, Aury J-M, Da Rocha M, Gouzy J, Sallet E, Martin-Jimenez C, Bailly-Bechet M, Castagnone-Sereno P, Flot J-F, et al: Hybridization and polyploidy enable genomic plasticity without sex in the most devastating plant-parasitic nematodes. PLoS Genet 2017, 13:e1006777.

77. Schiffer PH, Danchin E, Burnell AM, Schiffer A-M, Creevey C, Wong S, Dix I, O’Mahony G, Culleton BA, Rancurel C, et al: Signatures of the evolution of parthenogenesis and cryptobiosis in the genomes of panagrolaimid nematodes. bioRxiv 2017.

78. McLaren DJ: Oogenesis and fertilization in Dipetalonema viteae (Nematoda: Filarioidea). Parasitology 1973, 66:465–472.

79. Johnston WL, Krizus A, Dennis JW: Eggshell chitin and chitin-interacting proteins prevent polyspermy in C. elegans. Curr Biol 2010, 20:1932–1937.

80. Jaffe LA: Fast Block to Polyspermy in Sea-Urchin Eggs Is Electrically Mediated. Nature 1976, 261:68–71.

81. Wong JL, Wessel GM: Defending the Zygote: Search for the Ancestral Animal Block to Polyspermy. Current Topics in Developmental Biology 2005, 72:1–151.

82. Arnqvist Gr, Rowe L: Sexual conflict. Princeton, N.J.: Princeton University Press; 2005.

83. Birkhead TR, Pizzari T: Postcopulatory sexual selection. Nat Rev Genet 2002, 3:262–273.

84. Le Jambre LF, Royal WM: Meiotic abnormalities in backcross lines of hybrid Haemonchus. Int J Parasitol 1980, 10:281–286.

85. Chaudhry U, Redman EM, Abbas M, Muthusamy R, Ashraf K, Gilleard JS: Genetic evidence for hybridisation between Haemonchus contortus and Haemonchus placei in natural field populations and its implications for interspecies transmission of anthelmintic resistance. Int J Parasitol 2015, 45:149–159.

86. Great Britain. Ministry of Agriculture Fisheries and Food.: Manual of veterinary parasitological laboratory techniques. London,: H.M. Stationery Off.; 1971.

87. Denham DA: The development of Ostertagia circumcincta in lambs. J Helminthol 1969, 43:299–310.

88. Redmond DL, Smith SK, Halliday A, Smith WD, Jackson F, Knox DP, Matthews JB: An immunogenic cathepsin F secreted by the parasitic stages of Teladorsagia circumcincta. Int J Parasitol 2006, 36:277–286.

89. Kozarewa I, Ning Z, Quail MA, Sanders MJ, Berriman M, Turner DJ: Amplification-free Illumina sequencing-library preparation facilitates improved mapping and assembly of (G+C)-biased genomes. Nat Methods 2009, 6:291–295.

90. McKenna A, Hanna M, Banks E, Sivachenko A, Cibulskis K, Kernytsky A, Garimella K, Altshuler D, Gabriel S, Daly M, DePristo MA: The Genome Analysis Toolkit: a MapReduce framework for analyzing next-generation DNA sequencing data. Genome Res 2010, 20:1297–1303.

91. Danecek P, Auton A, Abecasis G, Albers CA, Banks E, DePristo MA, Handsaker RE, Lunter G, Marth GT, Sherry ST, et al: The variant call format and VCFtools. Bioinformatics 2011, 27:2156–2158.

92. Grattapaglia D, Sederoff R: Genetic linkage maps of Eucalyptus grandis and Eucalyptus urophylla using a pseudo-testcross: mapping strategy and RAPD markers. Genetics 1994, 137:1121–1137.

93. R Core Team: R: A Language and Environment for Statistical Computing. Vienna, Austria: R Foundation for Statistical Computing; 2015.

94. Broman KW, Wu H, Sen S, Churchill GA: R/qtl: QTL mapping in experimental crosses. Bioinformatics 2003, 19:889–890.

95. Broman KW: R/qtlcharts: interactive graphics for quantitative trait locus mapping. Genetics 2015, 199:359–361.

96. Zheng X, Levine D, Shen J, Gogarten SM, Laurie C, Weir BS: A high-performance computing toolset for relatedness and principal component analysis of SNP data. Bioinformatics 2012, 28:3326–3328.

